# Antigenic characterization of SARS-CoV-2 variants BA.3.2.1 and BA.3.2.2 in three animal models

**DOI:** 10.64898/2026.05.24.727525

**Authors:** Samuel A. Turner, Joey Olivier, Madison L. Ellis, Katharine A. Floyd, Lilin Lai, Suzanne M. Scheaffer, Ionsei Hastings, Tamarand L. Darling, Blake A. Miller, Charmy J. Patel, Hannah Peck, Daryll Vanover, Philip J. Santangelo, Michael S. Diamond, Mehul S. Suthar, Adrianus C.M. Boon, Derek J. Smith

**Author notes:** Correspondence to Mehul S. Suthar, Adrianus C.M. Boon, or Derek J. Smith. These authors contributed equally (Samuel A. Turner, Joey Olivier).

## Abstract

BA.3.2, a variant of SARS-CoV-2 containing ∼40 mutations in its spike protein compared to its nearest ancestor, has spread globally since its first detection in South Africa in November 2024. Here, we report antigenic characterization of BA.3.2 viruses in three naive animal models, and visualize its antigenic phenotype in the context of SARS-CoV-2 evolution using antigenic cartography. We find that: (1) BA.3.2 is substantially antigenically divergent from existing SARS-CoV-2 variants; (2) infection with BA.3.2 in hamster and mouse animal models produces sera with lower homologous titer than infection with other variants. Both of these results may have implications for the selection of vaccine antigens.

## Main text

BA.3.2, a highly mutated variant of SARS-CoV-2, was first detected in South Africa in November 2024^1,2^ and has since spread globally, accounting for ∼20% of SARS-CoV-2 sequences worldwide between April and May 2026^3^. Compared to its nearest ancestor, BA.3, the BA.3.2 spike carries 39 amino acid substitutions, two N-terminal domain deletions, and a four-residue insertion — meaning its emergence represents a saltation event similar to those which produced the early Omicron variants and BA.2.86^1,2^.

The large number of mutations in the spike protein of BA.3.2 suggests its antigenic phenotype may differ substantially from pre-existing variants.

Here, we report antigenic characterization of BA.3.2 using three naive animal models (see Supplementary methods for details):

1. Mouse serum following two doses of mRNA vaccine.
2. Mouse serum following infection.
3. Hamster serum following infection.

We characterize the two main phylogenetic clades of BA.3.2: BA.3.2.1, which is less prevalent and carries spike substitutions H681R and P1162R; and BA.3.2.2, which is more prevalent and carries spike substitutions K356T and A575S^3^.

We measured FRNT titers (see Supplementary methods for details) for BA.3.2 antigens against a panel of sera raised against historical and contemporary SARS-CoV-2 variants in the mouse mRNA vaccination model (both BA.3.2.1 and BA.3.2.2 antigens) and in the hamster infection model (BA.3.2.2 antigen only) (Figure 1). In the mouse mRNA vaccination model, fold-reduction from the homologous antigen to BA.3.2.1 and BA.3.2.2 antigens was largest for anti-KP.2 sera (>124x to >139x fold-reduction), and smaller both for early Omicron (anti-BA.1 and anti-BA.5) sera (>4.4x to >49x) and anti-XBB.1.5 sera (>10x to >20x). Where detectable, titers against the BA.3.2.1 antigen were nearly always lower than those to the BA.3.2.2 antigen (∼2-fold lower on average). In the hamster infection model, titers against the BA.3.2.2 antigen were non-detectable for nearly all sera (D614G, B.1.617.2, BA.1, BA.5, BA.2.86, JN.1, XFG).

**Figure 1.**
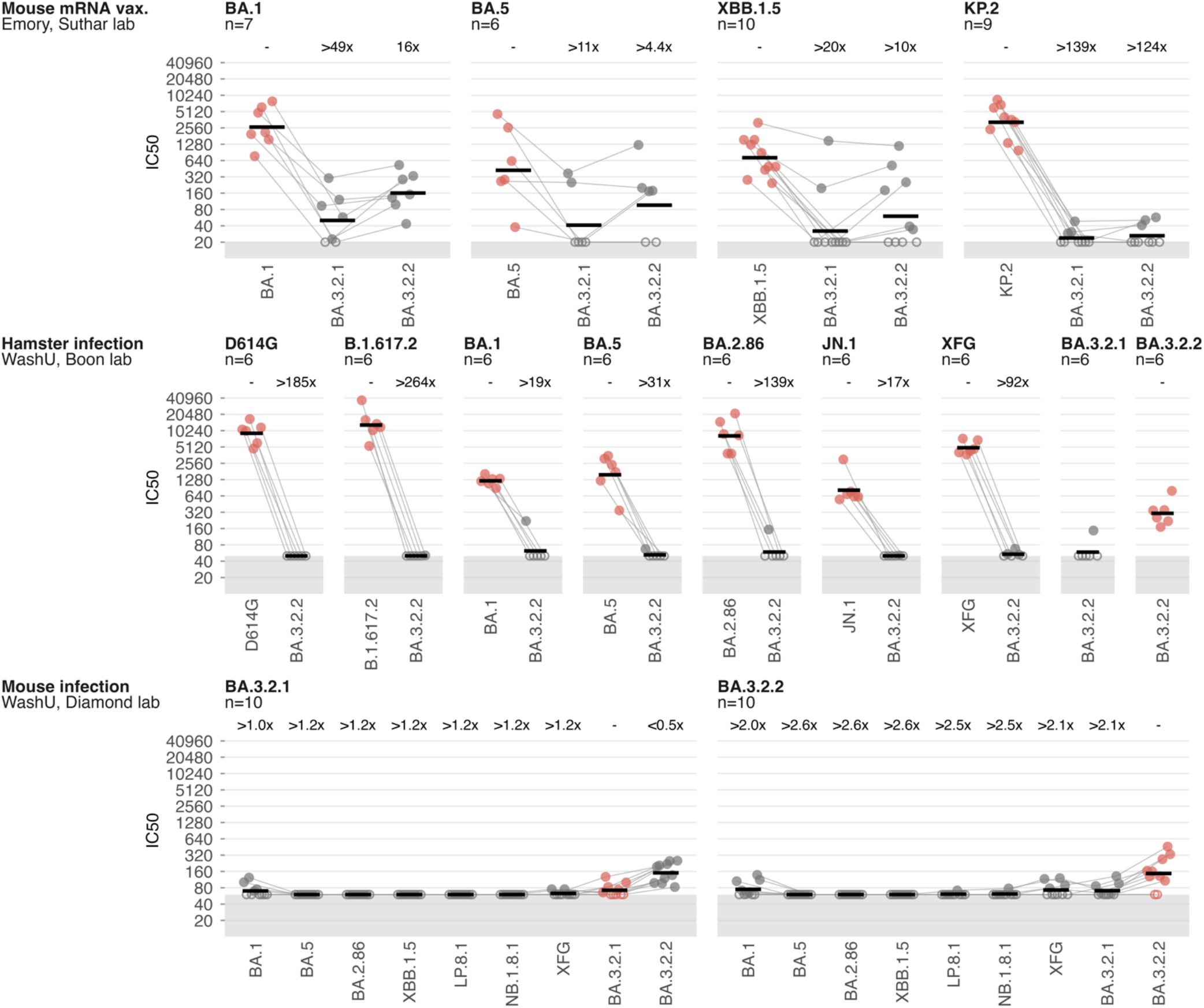
Neutralization titers against BA.3.2.1 and BA.3.2.2 antigens compared to historical and contemporary variants. Each panel shows sera raised against the antigen named in the panel header. Different rows show data from three different animal models. Red points show titers against the homologous antigen. Open circles show titers at or below the limit of detection (LOD). Black horizontal bars show the geometric mean titer, treating titers below the LOD as equal to the LOD. The geometric mean fold-reduction relative to the homologous antigen is written above each heterologous antigen. “>“ and “<“ indicate lower and upper bounds on the fold-reduction in cases where some titers were below the limit of detection.

In the hamster infection and mouse infection models, we raised sera against the BA.3.2.1 and BA.3.2.2 variants. In both models, homologous titers for these sera were substantially lower than for sera raised to other variants (Figure S1). In the hamster infection model, the majority of titers for anti-BA.3.2.1 sera against the BA.3.2.2 antigen were non-detectable. In contrast, in the mouse infection model, anti-BA.3.2.1 sera had a geometric mean titer of ∼160 to the BA.3.2.2 antigen, approximately 2-fold higher than their homologous titer to the BA.3.2.1 antigen. This latter observation indicates potentially higher avidity of the BA.3.2.2 antigen than the BA.3.2.1 antigen, which may also contribute to the titer differences between these antigens in the mouse mRNA vaccination data. The substantial majority of titers for anti-BA.3.2.1 and anti-BA.3.2.2 sera against other variants were non-detectable in the mouse infection model, although titers were detectable against BA.1 and XFG antigens more often than against other antigens.

We visualized the antigenic phenotype of the BA.3.2.1 and BA.3.2.2 variants in the context of historical SARS-CoV-2 variants using antigenic cartography (Figure 2)^4^. In the mouse mRNA vaccination map, BA.3.2.1 and BA.3.2.2 occupy novel antigenic space distinct from any pre-existing variant. Their antigenic positions are located further from JN.1-lineage antigens than from early Omicron (BA.1, BA.2, BA.5) and XBB.1.5 antigens, consistent with the larger fold-reduction observed for anti-KP.2 sera than for early Omicron and XBB.1.5 sera in this model. BA.3.2-lineage and JN.1-lineage antigens are approximately equidistant from pre-Omicron and early Omicron antigens. This distance matters because most individuals’ first SARS-CoV-2 exposures were to pre-Omicron or early Omicron variants — often in the form of highly immunogenic mRNA vaccines — resulting in substantial immune imprinting on those variants^5–8^. The comparable antigenic distances from these antigens to BA.3.2 and to JN.1-lineage variants are therefore consistent with observations in multiple human cohorts of similar neutralizing titer to BA.3.2-lineage and JN.1-lineage variants^9–11^.

**Figure 2.**
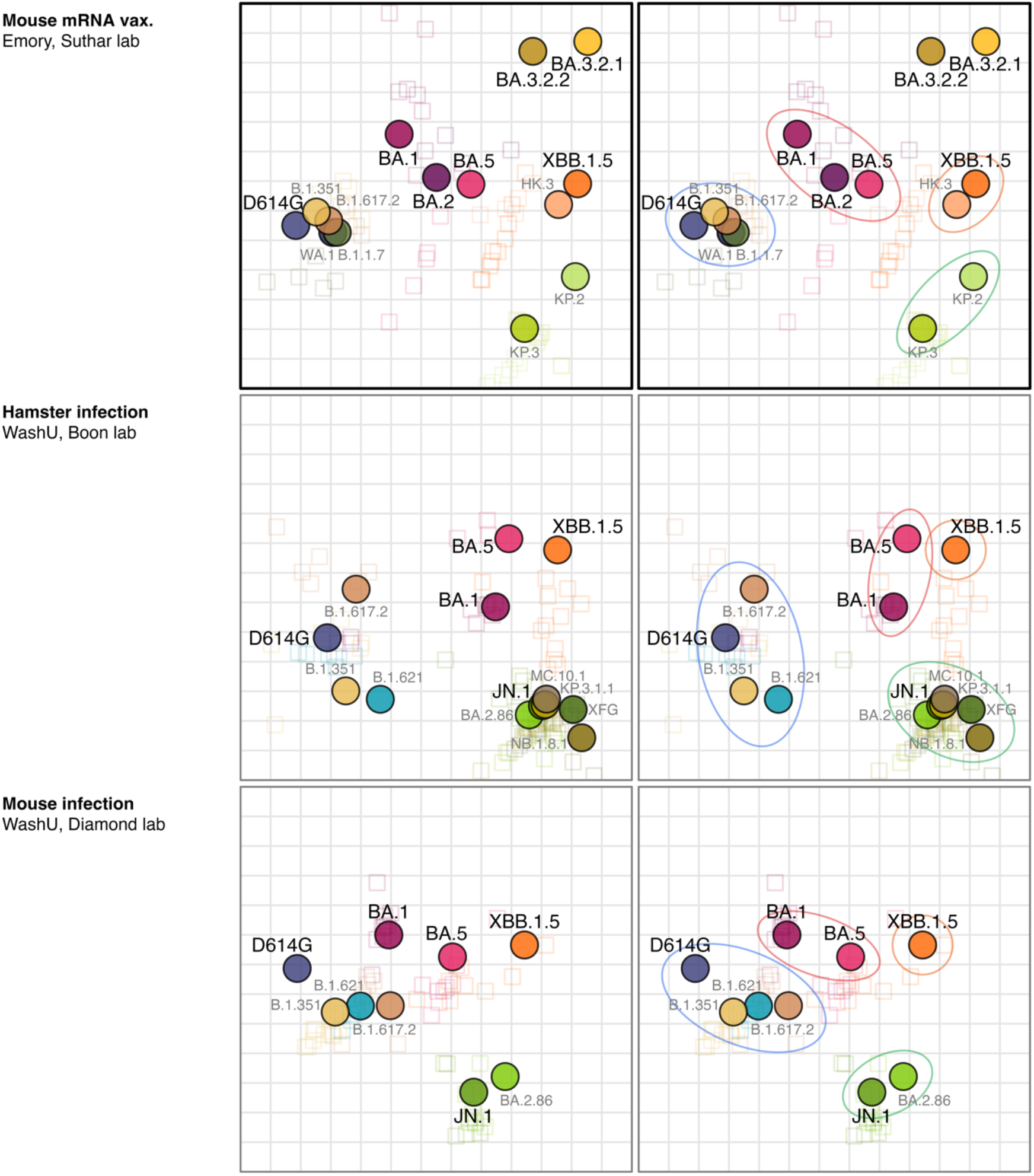
Antigenic maps for the three serum panels. Each row shows a map from one animal model, listed in the left column. Ellipses in the right-hand maps show the major antigenic groups: ancestral lineages, blue; early Omicron, red; XBB lineage, orange; JN.1 lineage, green. Points are antigens, colored by variant; open squares are individual sera. One grid square represents one antigenic unit, corresponding to a two-fold change in titer. All maps are Procrustes-aligned to the mRNA vaccination map (Emory, Suthar). BA.3.2.1 and BA.3.2.2 antigens and sera are not plotted in the hamster and mouse infection maps, as they could not be positioned due to the small number of detectable titers for these variants in these models.

Due to the small number of detectable titers for BA.3.2.1 and BA.3.2.2 antigens and sera in the hamster and mouse infection models, it was not possible to resolve the antigenic position of the variants in these maps.

These data indicate that BA.3.2 is substantially antigenically divergent from any existing SARS-CoV-2 variant. As such, despite the relatively modest reduction in human serum titers from the currently circulating JN.1-lineage variants (XFG and NB.1.8.1) to BA.3.2, BA.3.2 is nonetheless substantially antigenically distinct. This is consistent with observations that vaccination with a JN.1-lineage variant only modestly boosts BA.3.2 titers^9–11^, and suggests the boost to JN.1-lineage titers from a BA.3.2 vaccination may be similarly modest. The observation of low homologous titer for anti-BA.3.2 sera in the hamster and mouse infection models may also be relevant when considering the potential immunogenicity of a BA.3.2 vaccine antigen.

## Author contributions

Conceptualization: A.C.M.B., D.J.S., M.S.D., S.M.S., M.S.S., P.J.S.

Data curation: I.H., T.L.D., M.L.E., L.L.

Formal analysis: S.M.S., J.O., S.A.T., M.L.E., L.L.

Funding acquisition: A.C.M.B., D.J.S., M.S.S.

Investigation: J.O., I.H., T.L.D., S.M.S., S.A.T.

Methodology: I.H., T.L.D., M.S.D., S.M.S., S.A.T., J.O., B.A.M., C.J.P., H.P., D.V., K.A.F., M.L.E., L.L.

Project administration: A.C.M.B. Resources: A.C.M.B., P.J.S., M.S.S.

Software: J.O., S.A.T.

Supervision: A.C.M.B., D.J.S., M.S.S., P.J.S.

Visualization: S.A.T., J.O. Writing, original draft: S.A.T., J.O.

Writing, review & editing: A.C.M.B., M.S.S., D.J.S., S.A.T., J.O.

## Acknowledgements

We would like to thank members of the Boon-lab for technical and logistical support. We would also like to thank members of the NIH-SAVE team for providing isolates of SARS-CoV-2.

## Supplementary methods

### Neutralization assays

#### Mouse mRNA vaccination, Suthar laboratory

Live virus based focus reduction neutralization test (FRNT) assays were carried out following previously established protocols with minor adjustments^12–14^. Serum samples were tested in singlicates and subjected to eight serial three-fold dilutions in DMEM, starting at an initial dilution of 1:10. Each diluted serum sample was combined with an equal volume of live SARS-CoV-2 variant virus, corresponding to approximately 100 to 200 infectious foci per well. The serum virus mixtures were incubated for 1 hour at 37°C in round-bottom 96-well plates to allow antibody virus interactions. Following incubation, the mixtures were transferred onto monolayers of VeroE6-TMPRSS2 cells and incubated for an additional hour at 37°C. After this adsorption step, the inoculum was removed and replaced with 100 µl of prewarmed 0.85% methylcellulose overlay medium in each well to restrict viral spread. Plates were then incubated at 37°C for 18 to 40 hours depending on the replication kinetics of the specific viral variant. At the end of the incubation period, the overlay medium was carefully removed, and cells were washed with PBS and fixed using 2% paraformaldehyde for 30 minutes. Fixed cells were washed twice with PBS and subsequently permeabilized for at least 20 minutes using permeabilization buffer. Viral foci were detected by incubating the cells overnight at 4°C with an Alexa Fluor 647 conjugated anti-SARS-CoV-2 spike monoclonal antibody (CR3022-AF647). Plates were washed twice with PBS before imaging and quantification using an ELISPOT reader (CTL Analyzer).

#### Hamster infection, Boon laboratory

Serial dilutions of serum samples, starting at 1:50, were incubated with 10^2 focus-forming units of SARS-CoV-2 variants for 1 h at 37°C. Antibody-virus complexes were added to Vero-hTMPRSS2 cell monolayers in 96-well plates and incubated at 37°C for 1 hour. Subsequently, cells were overlaid with 1% (w/v) methylcellulose in Eagle’s minimal essential medium (MEM, Thermo Fisher Scientific) supplemented with 2% FBS. Plates were fixed 24 or 48 h later, dependent on the variant, with 10% formalin in PBS for 20 min at room temperature. Plates were washed and sequentially incubated with a pool of anti-S murine antibodies (SARS-2-02, -08, -09, -10, -11, -13, -14, -17, -20, -26, -27, -28, -31, -38, -41, -42, -44, -49, -57, -62, -64, -65, -67 and -71) plus anti-N of SARS-CoV-2, and HRP-conjugated goat anti-mouse IgG (Sigma-Aldrich catalog no. A8924) in PBS supplemented with 0.05% saponin and 2% FBS. SARS-CoV-2–infected cell foci were visualized using TrueBlue peroxidase substrate (KPL) and quantitated on an ImmunoSpot microanalyzer (Cellular Technologies).

#### Mouse infection, Diamond laboratory

FRNTs in the Diamond laboratory were performed as in the Boon laboratory, except serial dilutions of serum samples started at 1:60.

### mRNA formulation

mRNA constructs encoding pre-fusion stabilized Spike proteins were codon optimized and designed using human beta-globin 3’ UTR and cloned into a T7 driven expression pUC57 plasmid. Plasmids (GenScript, Piscataway, NJ) were linearized with NotI-HF (NEB, Ipswich, MA) overnight at 37°C, purified by sodium acetate (Thermo Fisher Scientific, Waltham, MA) precipitation and rehydrated with nuclease-free water. In vitro transcription was performed for 4 h at 37°C using the HiCap T7 kit (Aldevron, Fargo, ND) following the manufacturer’s instructions (N1-methyl-pseudouridine modified). The resulting RNA was treated with DNase I (Aldevron) for 30 min to remove the template, and it was purified using lithium chloride precipitation (Thermo Fisher Scientific). The RNA was heat-denatured at 65°C for 10 min before capping with a Cap-1 structure using guanylyl transferase and 2’-O-methyltransferase (Aldevron). The mRNA was then purified by lithium chloride precipitation, treated with alkaline phosphatase (NEB) and purified again. The mRNA concentration was measured using a NanoDrop instrument. Purified mRNA products were analyzed by capillary gel electrophoresis to ensure purity (Agilent 5200 Fragment Analyzer, Santa Clara, CA).

### Animal sera

#### Mouse mRNA vaccination, Suthar laboratory

Mouse studies were performed in strict accordance with the recommendations outlined in the Guide for the Care and Use of Laboratory Animals (National Institutes of Health). All protocols received approval from the Institutional Animal Care and Use Committee at Emory University (PROTO201700309). Female C57BL/6 mice (8-10 weeks; Jackson Labs) were vaccinated with 1 µg mRNA-LNP encoding prefusion-stabilized Spike protein through the intramuscular route. The second dose was administered four weeks later and serum was collected 30 days post-second dose. Serum was stored at -80°C and used for virus-neutralization assays.

#### Hamster infection, Boon laboratory

Animal studies were performed in strict accordance with the recommendations outlined in the Guide for the Care and Use of Laboratory Animals (National Institutes of Health). All protocols received approval from the Institutional Animal Care and Use Committee at Washington University School of Medicine (assurance number A3381-01). Male Syrian hamsters (5–6 weeks old) were procured from Charles River Laboratories or Inotiv and housed at WashU. The animals received 10^4 infectious units of D614G, B.1.351, B.1.621, B.1.617.2 or 10^5 infectious units of JN.1, BA.2.86, BA.1, BA.5, XBB.1.5, XFG, BA.3.2.1, BA.3.2.2 variants of SARS-CoV-2 (n = 5-6 hamsters per SARS-CoV-2 variant). Three weeks later, the animals were sacrificed and blood was collected via cardiac puncture. The collected blood was stored at 4°C overnight. Serum was collected after centrifugation of the clotted blood for 20 minutes. Serum was stored at 4°C and used for virus-neutralization assays.

#### Mouse infection, Diamond laboratory

8-week-old female K18-hACE2 C57BL/6J mice (strain: 2B6.Cg-Tg(K18-ACE2)2Prlmn/J, Cat # 34860, Jackson Laboratory) were inoculated intranasally after anesthesia with xylazine and ketamine hydrochloride with: 10^4 FFU of BA.1, BA.5, BA.2.86, JN.1, BA.3.2.1, or BA.3.2.2; 10^3 FFU of B.1.621, or XBB.1.5; or 10^1 FFU of B.1.617.2 or D614G. B.1.351 sera were produced at both 10^2 and 10^3 FFU doses. At 35 days post-infection, serum was collected and mice were administered terminal anesthesia with a ketamine overdose.

### Antigenic cartography

Antigenic cartography allows quantification of antigenic differences between variants and strains. Briefly, each antigen-serum titer is transformed into a target distance between the antigen and serum, by calculating the log2 fold-reduction from the highest titer observed for the serum. Coordinates for each antigen and serum are then optimized to minimize the discrepancy between the target antigen-serum distances and those in the map. A detailed description is available in Smith et al. 2004^4^ and in the documentation of the Racmacs package^15^.

Antigenic cartography was performed using the Racmacs package (v1.2.9) in R (v4.5.1). Each map was constructed using 1000 optimizations, with a dilution step size of 0, and with the minimum column basis parameter set to “none”.

When constructing antigenic maps, we:

1. perform a few types of adjustment to titers to account for various features of titer data which do not relate to the underlying antigenic phenotype;
2. impute some titers, when doing so resolves substantial mispositioning of variants caused by missing data that can be sensibly estimated.

We first describe the types of adjustment or titer imputation, then describe the specific applications to each map. Adjustments and imputations apply only to the maps in Figure 2; titer values in Figures 1 and S1 are unadjusted.

#### Adjusting for reactivity differences between titration sets

Two separate titration sets sometimes contain the same antigen titrated against overlapping but non-identical sets of sera, with the antigen showing systematically higher titers in one set than the other for the overlapping sera. In these cases, we correct for the reactivity differences as follows:

1. Choose a titration set to use as the reference set; designate the other the “target” titration set.
2. Calculate the mean log_2_ fold-change from the reference titration set to the target titration set for sera shared between the two.
3. Discard the titrations of the antigen against the shared sera from the target titration set.
4. Subtract the mean log_2_ fold-change from step 2 from the titers in the target titration set against non-shared sera.

#### Adjusting for antigen avidity

Antigens vary in their baseline assay reactivity (“avidity”), independent of their antigenic similarity to the sera they are titrated against^16^. For example, if an antigen produces a titer against a serum that exceeds the serum’s titer against its own homologous antigen — and the antigen does so for multiple sera — it is likely to be a high-avidity antigen. We correct for avidity by applying a manually chosen log-scale shift to every titer against the antigen, using the `optimizeAgReactivity` function in Racmacs.

#### Adjusting for non-specific inhibition

Some antigens produce a non-specific signal meaning that titers do not fall below a baseline value, even against distantly related sera. For these antigens, we treat values at or near this baseline as upper bounds on the titer (e.g., titer 100 becomes <100).

#### Adding reconstructed titers

Some sera in our dataset are titrated only against recent antigens, meaning they cannot be well-positioned relative to earlier variants. Where a distinct set of sera raised against the same variant was titrated against the missing antigens, we use those measurements to reconstruct the missing titers.

Let the reference titer table contain sera against variant V titrated against a broad range of antigens, and the target titer table contain distinct sera raised against V which were titrated against only a few recent antigens. For each serum s in the target titer table and each antigen A present in the reference table but not in the target table, we calculate:

GMFR(V to A), the geometric mean fold-reduction from V to A across all sera in the reference table raised against V.

We use this to set the titers for each serum s in the target table against antigens A from the reference table by assuming the V to A fold-reduction is the same on average across the two serum sets:

titer(s, A) = titer(s, V) / GMFR(V to A)

We do not modify measured titers in the target table.

#### Adding thresholded titers

Sometimes, a recent antigen has not been titrated against sera raised against early-pandemic variants. In these cases, there are no measured titers (and therefore target distances) constraining the antigen from being placed near to the early-pandemic variants. To resolve these cases, we identify a closely related recent ancestor (for example, for KP.3.1.1, we may choose JN.1) which was titrated against sera raised against early variants, and set the descendant antigen’s titer against early-pandemic sera to ≤ the corresponding titer of the chosen ancestor.

#### Mouse mRNA vaccination map, Suthar laboratory, Emory

We adjusted for antigen avidity for the KP.2 antigen by decreasing its reactivity by 2.5 log2 units. We reconstructed titers for BA.1, BA.5, XBB.1.5, and KP.2 sera for non-homologous antigens in titer table B (Table S1), using titer table A as the reference table.

#### Mouse infection map, Diamond laboratory, WashU

We adjusted for reactivity differences for the B.1.617.2 antigen between titer table A and titer table B (Table S2), taking titer table B as the reference table and titer table A as the target table. We adjusted for non-specific inhibition for BA.2.86 and JN.1 antigens’ titers against pre-XBB.1.5 sera. The NB.1.8.1 and LP.8.1 antigens were titrated but their position in the antigenic map was underconstrained so they could not be placed.

#### Hamster infection map, Boon laboratory, WashU

BA.1 sera from titer table A were excluded (Table S3). We adjusted for reactivity differences for B.1.351 and B.1.617.2 antigens between titer table A and titer table B, taking titer table B as the reference table and titer table A as the target table. We added thresholded titers for JN.1 derivatives KP.3.1.1, MC.10.1, NB.1.8.1, and XFG, using JN.1 as the closely related recent ancestor. Finally, we adjusted for antigen avidity for BA.1 (+1 log2 unit) and BA.5 (-1 log2 unit).

## Supplementary figures

**Figure S1.**
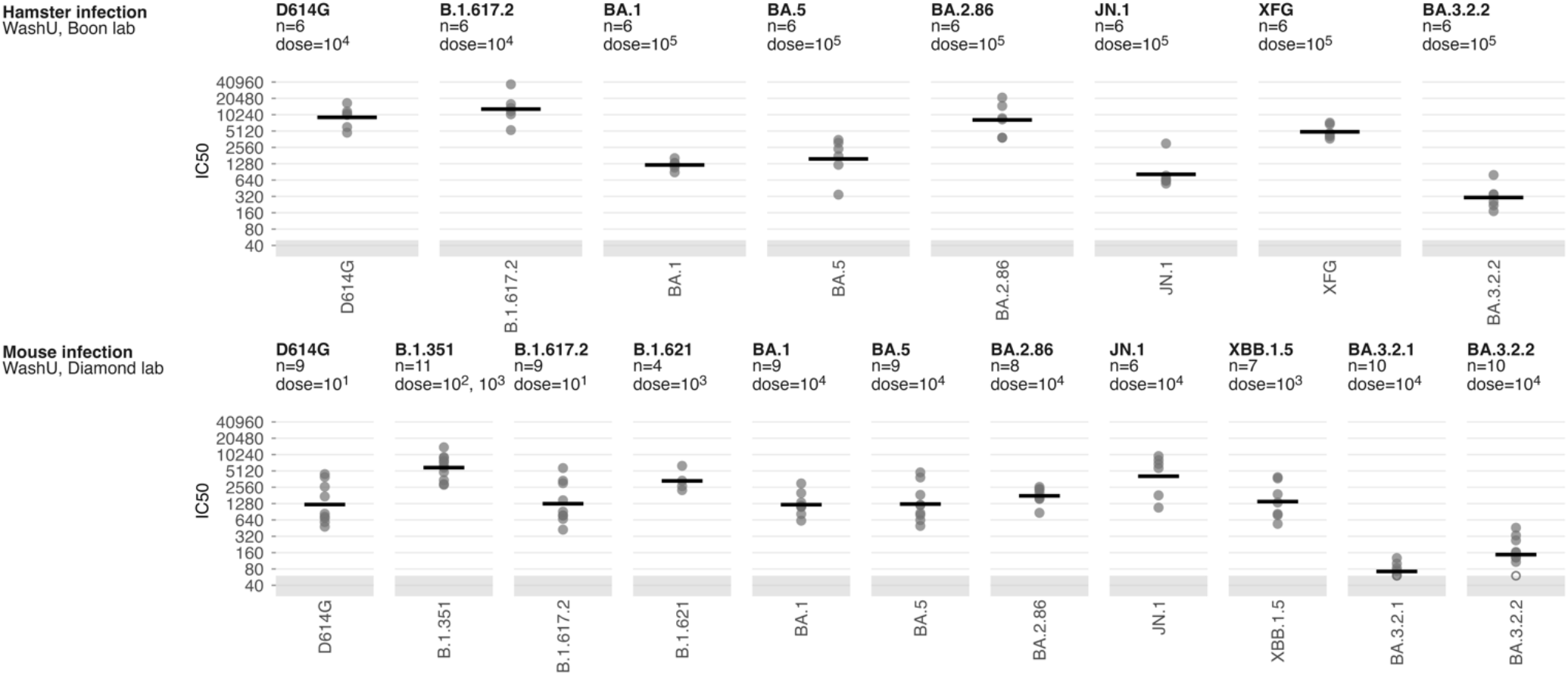
Homologous neutralization titers for the hamster infection and mouse infection serum panels. Each panel shows IC50 neutralization titers for a group of sera against its homologous antigen, with the number of sera and the inoculation dose given in the panel subtitle. Open circles show titers at or below the limit of detection (grey shaded band). Black horizontal bars show the geometric mean titer.

**Table S1.** Mouse vaccination titer tables, Suthar laboratory, Emory.

| Titer table A |  | Titer table B |  |
| --- | --- | --- | --- |
| Sera | Antigens | Sera | Antigens |
| WA.1 | WA.1 |  |  |
| D614G | D614G |  |  |
| B.1.1.7 | B.1.1.7 |  |  |
| B.1.617.2 | B.1.617.2 |  |  |
| B.1.351 | B.1.351 |  |  |
| BA.1 | BA.1 | BA.1 | BA.1 |
| BA.2 | BA.2 |  |  |
| BA.5 | BA.5 | BA.5 | BA.5 |
| XBB.1.5 | XBB.1.5 | XBB.1.5 | XBB.1.5 |
| HK.3 | HK.3 |  |  |
| KP.2 | KP.2 | KP.2 | KP.2 |
| KP.3 | KP.3 |  |  |
|  |  |  | BA.3.2.1 |
|  |  |  | BA.3.2.2 |

**Table S2.** Mouse infection titer tables, Diamond laboratory, WashU.

| Titer table A |  | Titer table B |  | Titer table C |  |
| --- | --- | --- | --- | --- | --- |
| Sera | Antigens | Sera | Antigens | Sera | Antigens |
| D614G | D614G | D614G | D614G |  |  |
| B.1.617.2 | B.1.617.2 | B.1.617.2 | B.1.617.2 |  |  |
| B.1.351 | B.1.351 | B.1.351 | B.1.351 |  |  |
| B.1.621 | B.1.621 |  |  |  |  |
| BA.1 | BA.1 |  |  |  |  |
| BA.5 | BA.5 |  |  |  |  |
| XBB.1.5 | XBB.1.5 |  |  |  | XBB.1.5 |
| BA.2.86 | BA.2.86 |  |  |  | BA.2.86 |
| JN.1 | JN.1 |  |  |  |  |
|  |  |  |  |  | LP.8.1 |
|  |  |  |  |  | XFG |
|  |  |  |  |  | NB.1.8.1 |
|  |  |  |  | BA.3.2.1 | BA.3.2.1 |
|  |  |  |  | BA.3.2.2 | BA.3.2.2 |

**Table S3.**
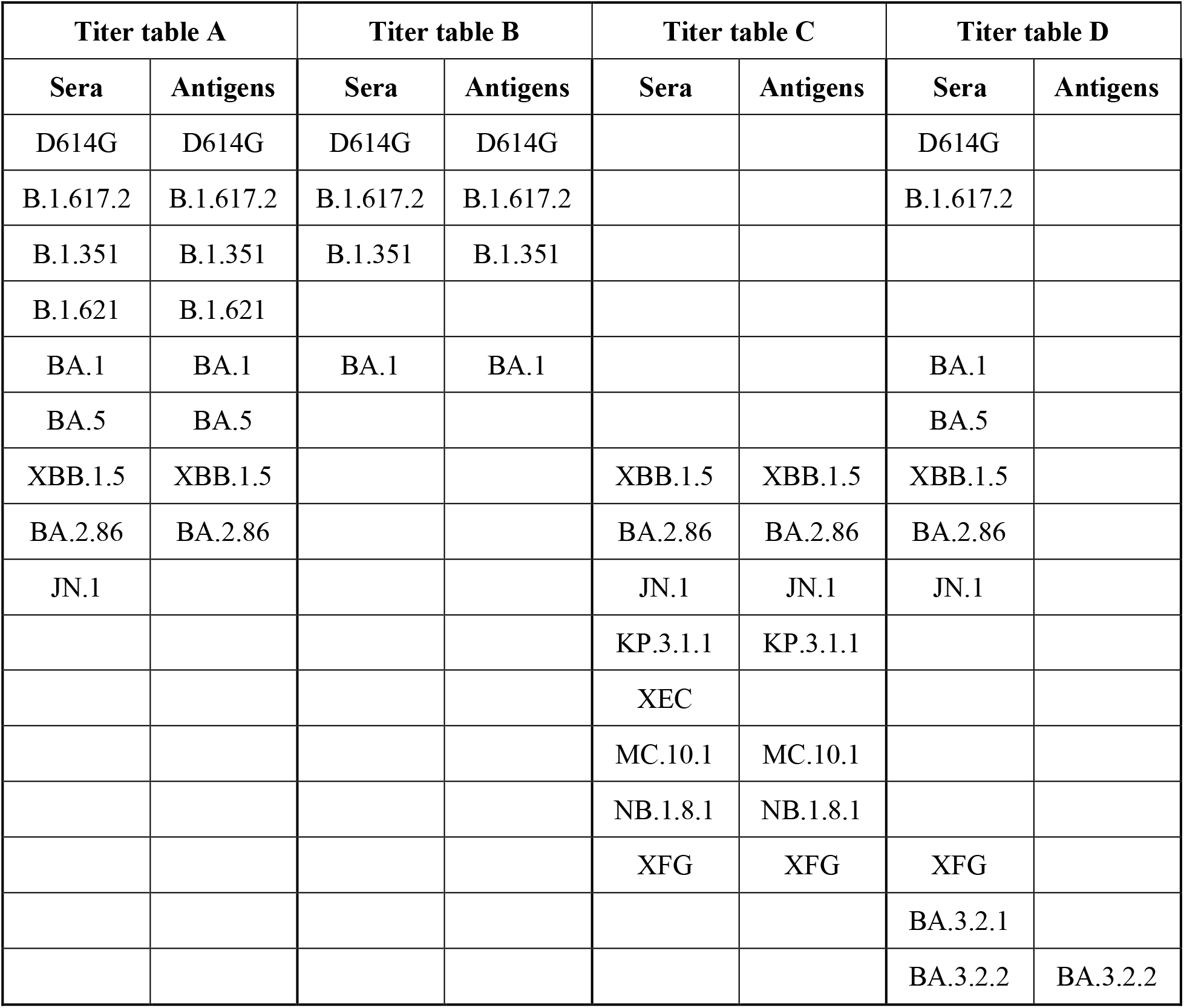
Hamster infection titer tables, Boon laboratory, WashU.

| Titer table A |  | Titer table B |  | Titer table C |  | Titer table D |  |
| --- | --- | --- | --- | --- | --- | --- | --- |
| Sera | Antigens | Sera | Antigens | Sera | Antigens | Sera | Antigens |
| D614G | D614G | D614G | D614G |  |  | D614G |  |
| B.1.617.2 | B.1.617.2 | B.1.617.2 | B.1.617.2 |  |  | B.1.617.2 |  |
| B.1.351 | B.1.351 | B.1.351 | B.1.351 |  |  |  |  |
| B.1.621 | B.1.621 |  |  |  |  |  |  |
| BA.1 | BA.1 | BA.1 | BA.1 |  |  | BA.1 |  |
| BA.5 | BA.5 |  |  |  |  | BA.5 |  |
| XBB.1.5 | XBB.1.5 |  |  | XBB.1.5 | XBB.1.5 | XBB.1.5 |  |
| BA.2.86 | BA.2.86 |  |  | BA.2.86 | BA.2.86 | BA.2.86 |  |
| JN.1 |  |  |  | JN.1 | JN.1 | JN.1 |  |
|  |  |  |  | KP.3.1.1 | KP.3.1.1 |  |  |
|  |  |  |  | XEC |  |  |  |
|  |  |  |  | MC.10.1 | MC.10.1 |  |  |
|  |  |  |  | NB.1.8.1 | NB.1.8.1 |  |  |
|  |  |  |  | XFG | XFG | XFG |  |
|  |  |  |  |  |  | BA.3.2.1 |  |
|  |  |  |  |  |  | BA.3.2.2 | BA.3.2.2 |

